# Semiology, clustering, periodicity and natural history of seizures in an experimental visual cortical epilepsy model

**DOI:** 10.1101/289256

**Authors:** Bao-Luen Chang, Leite Marco, Albert Snowball, Elodie Chabrol, Andreas Lieb, Matthew C. Walker, Dimitri M. Kullmann, Stephanie Schorge, Robert C. Wykes

**Affiliations:** Department of Clinical and Experimental Epilepsy, UCL Institute of Neurology, Queen Square, London WC1N 3BG, UK; Section of Epilepsy, Department of Neurology, Chang Gung Memorial Hospital Linkou Medical Center and Chang Gung University College of Medicine, Taoyuan, Taiwan

**Keywords:** Tetanus toxin, Visual cortical epilepsy, Circadian rhythm, Seizure clustering, Periodic pattern, Prediction

## Abstract

**Objective:** To characterize a rat model of focal neocortical epilepsy for use in developing novel therapeutic strategies in a type of epilepsy that represents a significant unmet need.

**Methods:** Intracortical tetanus toxin (TeNT) injection was used to induce epilepsy in rats. Seizures and their behavioural manifestations were evaluated with continuous video-electrocorticography telemetry.

**Results:** TeNT injection into rat primary visual cortex induced focal neocortical epilepsy without preceding status epilepticus. The latency to first seizure ranged from 3 to 7 days. Seizure duration was bimodal, with both short (approximately 30s) and long-lasting (>100s) seizures occurring in the same animals. Seizures were accompanied by non-motor features such as behavioural arrest, or motor seizures with or without evolution to generalized tonic-clonic seizures. Seizures were commoner during the sleep phase of a light-dark cycle. Seizure occurrence was not random, and tended to cluster with significantly higher probability of recurrence within 24 hours of a previous seizure. Across animals, the number of seizures in the first week could be used to predict the number of seizures in the following 22 days.

**Significance:** The TeNT model of visual cortical epilepsy is a robust model of acquired focal neocortical epilepsy, and is well suited for preclinical evaluation of novel anti-epileptic strategies. We provide here a detailed analysis of the epilepsy phenotype, seizure activity, electrographic features, and the semiology. In addition we provide a predictive framework that can be used to reduce variation and consequently animal use in pre-clinical studies of potential treatments.

**Key Points:**

- Tetanus toxin injection into rat visual cortex induces focal cortical epilepsy.
- Electrographic seizures were associated with non-motor and motor features with or without evolution to generalized tonic-clonic seizures.
- Seizures could not be provoked by intermittent photic stimulation.
- Seizures were clustered in time and exhibited a circadian variation in frequency.
- The number of seizures in first week after seizure onset could be used to predict the total number of seizures in the following 3 weeks.

## Introduction

Animal models of epilepsies are an invaluable tool to test novel anti-epileptic strategies. At present, the most commonly used rodent models of epilepsy rely on chemoconvulsants such as kainic acid or pilocarpine, or electrical stimulation, to induce status epilepticus, which is followed by spontaneous recurrent seizures. These predominantly model mesial temporal epilepsy (mTLE).^1^ Various genetic mouse and rat models have also been used, mainly to model primary generalized epilepsy. In utero electroporation can also be used to manipulate phosphatase and tensin homolog (Pten) or related signalling molecules to generate malformations of cortical development.^2^ Although all these models are useful, there is a need for a robust model of adult-onset neocortical epilepsy with a well-defined focus. Focal epilepsy has been estimated to account for ∼60% of human cases. It is frequently pharmacoresistant,^3, 4^ and therefore represents a major unmet need. It is therefore necessary to develop an optimal animal model of focal neocortical epilepsy for epilepsy research.

Tetanus neurotoxin (TeNT) is a metalloprotease (zinc-dependent protease) that specifically targets neurons.^5^ TeNT consists of a heavy chain and a light chain with a disulphide bond linked between both chains. Intact tetanus neurotoxin is necessary for uptake by synaptic terminals since the function of the heavy chain is for specific binding and internalization into neurons, as well as retrograde axonal transport.^6^ The action on neurotransmitter release depends on the light chain, and requires intracellular cleavage of the inter-chain disulphide bond.^6^ The activated light chain specifically cleaves synaptobrevin which is a vesicle-associated membrane protein (VAMP) and a component of the SNARE (SNAP REceptor) complex that is critically responsible for presynaptic neurotransmitter release.^7, 8^ As a result, TeNT causes vesicles to fail to fuse with the presynaptic membrane disrupting exocytosis and the release of neurotransmitters. Furthermore, TeNT predominantly interferes with presynaptic vesicle release of inhibitory neurotransmitters, both γ-aminobutyric acid (GABA) and glycine, in CNS, and therefore disinhibits the local circuitry.^9^

Injection of TeNT into hippocampus has been used to produce a rat model of mesial temporal lobe epilepsy.^10, 11^ A rat model which may share similarities to human epilepsia partialis continua (EPC) has been induced by TeNT injection to motor cortex.^12^ However, the major feature of this model is brief bursts of high frequency (<1 s, 120 – 160 Hz) EEG activity. Although discrete long-lasting epileptic discharges and generalized motor convulsions can be elicited with higher doses of TeNT, this is relatively poorly tolerated with a high proportion of animals experiencing >15% weight loss or death following a severe seizure.^12, 13^ A mouse model using TeNT in visual cortex has been reported. However, this elicited electrographic epileptiform discharges with clusters of high-amplitude spikes lasting for >4 s, and behavioural correlates were not reported.^14^

The goal of this study is to characterize the rodent visual cortex TeNT model to study focal neocortical epilepsy, suitable for future translation of experimental treatments. We comprehensively characterize the epilepsy phenotype, seizure activity, electrographic features, and the semiology. We identified a diurnal fluctuation in seizure frequency, as well as non-random clustering and periodic phenomena. Finally, we further provide a prediction profile on seizure activity in this TeNT model of visual cortical epilepsy which can contribute to future studies using this model.

## Methods

### Animals

All animal experiments were conducted in accordance with the United Kingdom Animal (Scientific Procedures) Act 1986, and approved by the local ethics committee. Male Sprague-Dawley rats (6 – 12 weeks old, 260 – 330 g; Charles River, UK) were used for all the analysis, except Lister Hooded and Long-Evans rats were used to validate the model in additional strains and for the intermittent photic stimulation. Animals were housed on a 12/12 hour light/dark cycle (light cycle 7 a.m./7 p.m.), and maintained under controlled environmental conditions at ambient temperature 24-25 °C and humidity 50-60 % with environmental enrichment and *ad libitum* access to food and water. Animals were group housed and allowed to acclimatize to the new environment for at least 1 week before surgery and housed individually after surgery.

### Stereotactic surgery and implantation of wireless ECoG telemetry system

Rats were anaesthetized using isoflurane and placed in a stereotaxic frame (Kopf, USA) with head-fixed. 15 – 15.6 ng of tetanus toxin (gift of G. Schiavo) adjusted with animal’s body weight in a final volume of 1 µl was injected into layer 5 of right primary visual cortex at a rate of 200 nl/min (coordinates: 3 mm lateral and 7 mm posterior of bregma at a depth of 1 mm from pia). An electrocorticography (ECoG) transmitter (A3028E-AA, Open Source Instruments^15^, MA, USA) was implanted subcutaneously with a recording electrode positioned on the visual cortex, and a reference electrode was placed in the contralateral frontoparietal cortex. Animals were single housed in Faraday cages and 24 h/7 days telemetric ECoG was continuously recorded post surgery and up to 30 days after the onset of first spontaneous seizure for the individual animals. For the rats in vehicle control group, all the same surgical procedures were performed except 1 µl of 0.9% saline was injected in right primary visual cortex instead of tetanus toxin.

### Video-ECoG monitoring, ECoG data acquisition and analysis

IP cameras (M7D12POE or M7D12HD, Microseven) with infrared video surveillance system and time-locked to the ECoG were used for continuous 24 h/7 days telemetric video-ECoG monitoring. ECoG was recorded and processed using the Neuroarchiver tool (Open Source Instruments). The sampling rate of A3028E-AA implantable transmitters is 512 SPS (sample per second) which provides the frequency dynamic range (bandwidth) of 0.3 – 160 Hz and voltage dynamic range is approximately 20 mV (-13 mV to +7 mV). Artefacts consisting of short duration (< 100 ms), high amplitude “glitches” were excluded. The ECoG was interpreted and epileptic seizures were detected manually by a neurologist continuously analyzing the entire ECoG dataset. The criteria used for detecting epileptic seizures are an evolution of frequency and amplitude over time with a sudden, repetitive, rhythmic, evolving and stereotypic abnormal electrographic activity with high amplitude (>2 times that of baseline) and a minimum duration of 10 seconds.^16–19^

### Behavioural seizure analysis and classification

To establish the classification of seizure behaviours corresponding to human seizure semiology, the top-down video footage time-locked to the ECoG trace was used to assess the behavioural correlates of epileptic seizures. A subset of seizures was randomly selected from animals with telemetric video-ECoG recordings for up to 5 weeks from TeNT injection. Seizure types were evaluated and classified by a neurologist according to the ILAE 2017 classification of seizure types.^20^

### Intermittent photic stimulation

Intermittent flickering light stimulation was performed in Lister Hooded epileptic rats at around two weeks after the onset of spontaneous seizures. To enhance the effect of photic stimulation, all other lights in the room were switched off during the test. An LED strip with high light intensity (> 100 thousand foot-candles) attached to a board was wrapped around the cage. The frequency range of flickering light stimulation was from 1 Hz to 30 Hz. Different frequencies of photic stimulation were applied in the following sequence: 1Hz, 3Hz, 6Hz, 9Hz, 12Hz, 15Hz, 18Hz, 21Hz, 24Hz, 27Hz, and 30Hz, then followed by the reverse sequence of stimulation frequencies (from 30 Hz to 1 Hz). The duration of stimulation in each frequency was 20 seconds then rest for 20 seconds before starting the next sequential frequency of stimulation.

### Statistical Analysis

Data and statistical analyses were carried out as appropriate using Prism 6 (GraphPad) or Python (version 3.6, Anaconda, Inc. Texas, USA). To test whether the occurrence of seizures followed a Poisson process, the distributions of inter-seizure-intervals (ISIs) were compared to an exponential distribution with unknown mean using the Lilliefors adaptation to the Komolgorov-Smirnov test.^21^

The autocorrelation was calculated to understand if the seizures tend to arise in clusters. The peri-seizure histograms were computed by counting the intervals between all seizures and correcting for the fact that long intervals are less common due to the limited duration of the experiment of 30 days, i.e., each bin count was divided by 30 minus its central bin value in days. Two-tailed one sample t-tests were used to test if the mean of the histogram bins across rats (normalized by total seizure count) differed significantly from the counts expected from a uniform distribution.

The partial autocorrelation of daily seizure occurrence was computed to test the dynamical properties of seizure occurrence. Two-tailed one sample t-tests were used to test if the partial auto-correlation coefficients significantly differed from zero across rats.

A Gaussian process model using a linear kernel was fitted to the data in order to analyse the predictive power that the number of seizures in the first week of recording has upon the number of seizures occurring in the remaining days of the experiment.

## Results

### Tetanus toxin injection into visual cortex produces acquired occipital lobe epilepsy in rats

We developed and optimized a rat model of visual cortical epilepsy by administration of tetanus toxin into primary visual cortex of different rat strains, which produced clear, robust and long-lasting spontaneous seizures in all three strains tested (Figure 1 & Fig. S1). The electrocorticogram at seizure onset was often characterized by low-amplitude fast activity, which evolved in amplitude and frequency. This was followed by repetitive spikes, sharp waves, polyspikes or polyspikes-and-waves, which then slowed in frequency and there was EEG suppression at the end of a seizure (Figure 1A,B). The overall success rate of TeNT-elicited epilepsy in the visual cortex was approximately 84%. The TeNT treatment and seizures were well-tolerated by most animals. SUDEP (sudden unexpected death in epilepsy) or death due to the effects of TeNT were not observed. Animals that received TeNT injections but did not develop seizures, produced only epileptiform spikes. No seizures, epileptiform activity or behavioural abnormalities were observed in the vehicle control animals (n = 9). To reduce variability and increase the ability to compare with previous work in Sprague-Dawley rats, all the further detailed analyses were performed using Sprague-Dawley rats only.

**Figure 1.**
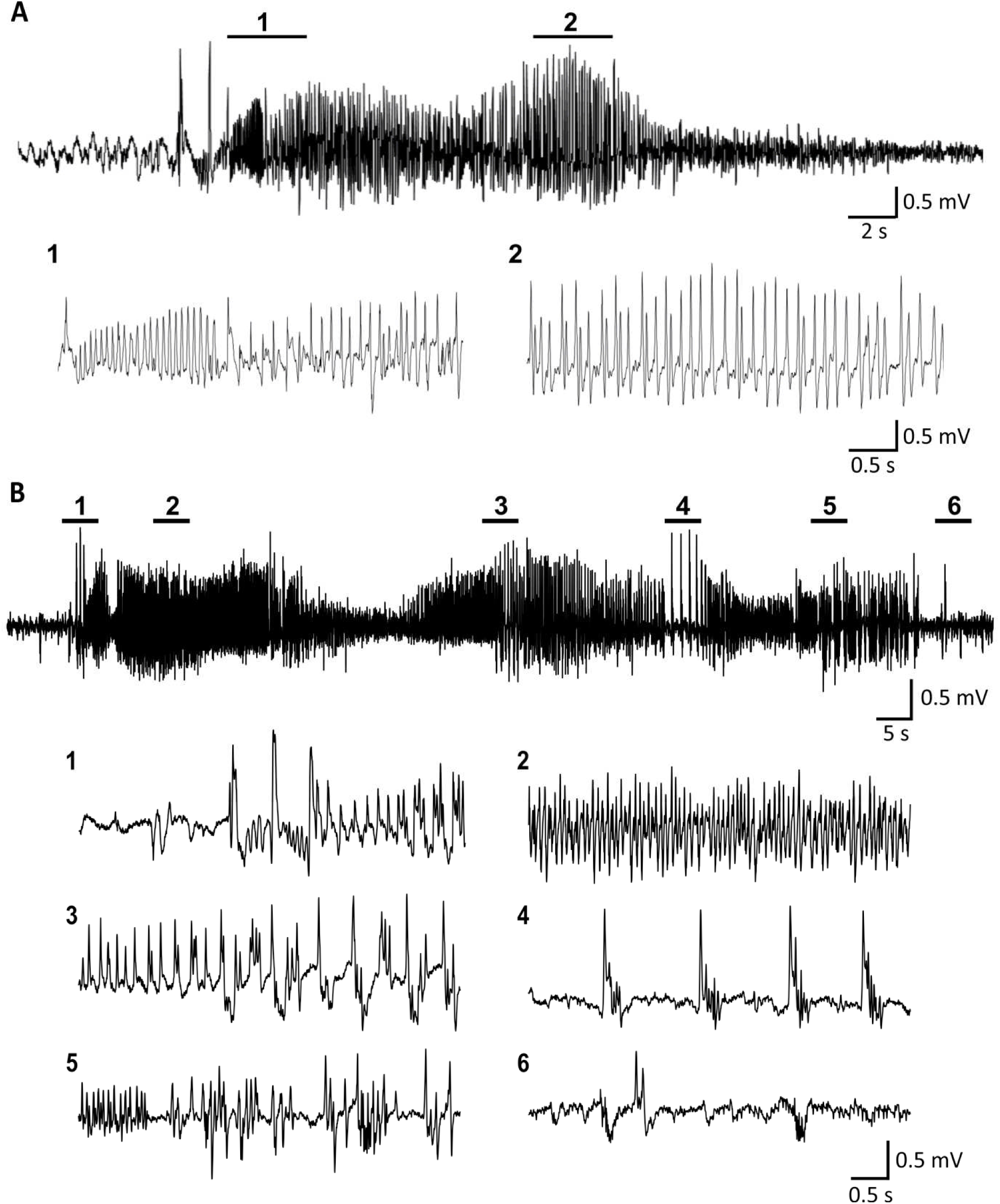
Representative ECoG features of ictal discharges from Sprague Dawley rats. A, a short seizure lasting < 40 s and B, a long-lasting seizure (> 100 s). The seizures start with fast activity evolving to high-amplitude and high-frequency waves with propagation followed by gradually subsided at the end of seizures.

The latent period, defined as the delay before the emergence of the first seizure after TeNT injection, ranged from 3 to 7 days (Figure 2A). The overall trend was for seizure frequency to gradually increase over time reaching a plateau 16 – 19 days after onset of seizures and decreasing thereafter in most animals (Figure 2B, left panel). As the total number of seizures experienced among the individual animals was highly variable ranging from tens to hundreds of seizures (Figure 3C, left panel), we normalised to the % of total seizures during different time periods in each individual animal to compare frequency patterns between animals. The average proportion of seizures was stable in the first 2 weeks (26% in 1st week; 24% in 2nd week), and increased to a peak in the 3rd week (36%) and then dropped to the lowest proportion (12%) in the last week of recordings (Figure 2B, right panel). The cumulative distribution of seizures in most of the animals followed a similar pattern showing a steady increase in the cumulative seizure number from the onset of seizures to 20 days after seizure onset followed by a plateau in the final days of recordings. Two animals had seizure activity that dramatically increased 15 – 20 days after the onset of first seizure (Figure 2C, right panel). In all animals, the seizure duration evolved over time as epileptic activity became established. The seizure duration was relatively short in the first few days after the onset of seizures (< 50 secs), but the duration gradually increased to a stable duration of approximately 100-120 secs from one week after the onset of seizures (Figure 2D).

**Figure 2.**
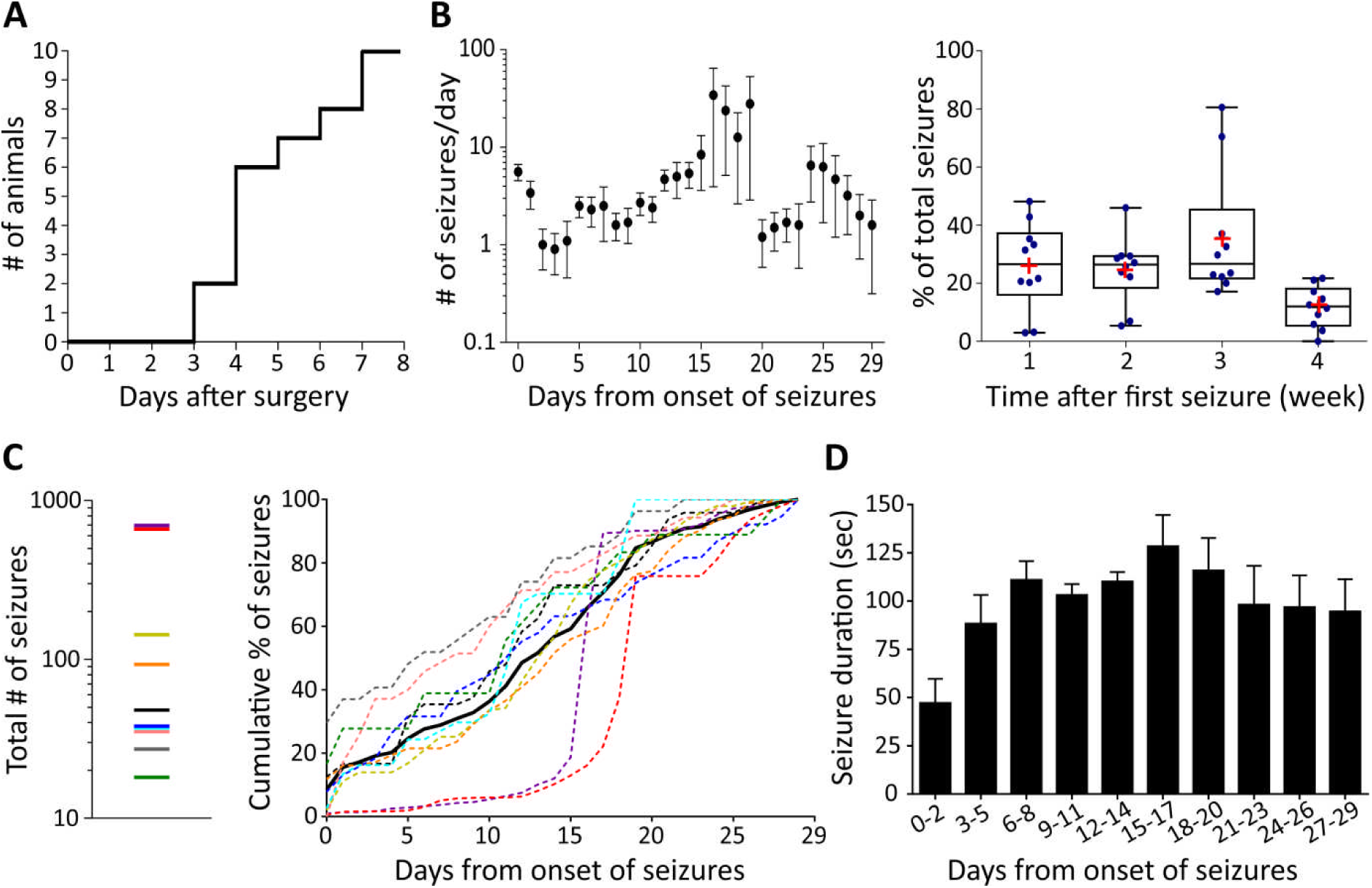
Characterization of the TeNT model of visual cortical epilepsy in rats. A, The onset of seizures after surgery was between 3 to 7 days and most were around 4 days. B, left: Represents the distribution of daily seizure frequency (mean ± SEM) during the whole recording period from the onset of 1st seizure (time point 0). Right, the box-and-whisker graph shows the mean (red cross) and the median of % of weekly seizure frequency. C, Total number of seizures (left panel) and the corresponding cumulative % distribution for individual animals (color dotted line) showing in right panel. The black line represents the mean of the values. D, The average of median of seizure duration from the onset of 1st seizure (time point 0). Data are presented as mean ± SEM. (n=10 Sprague-Dawley rats)

**Figure 3.**
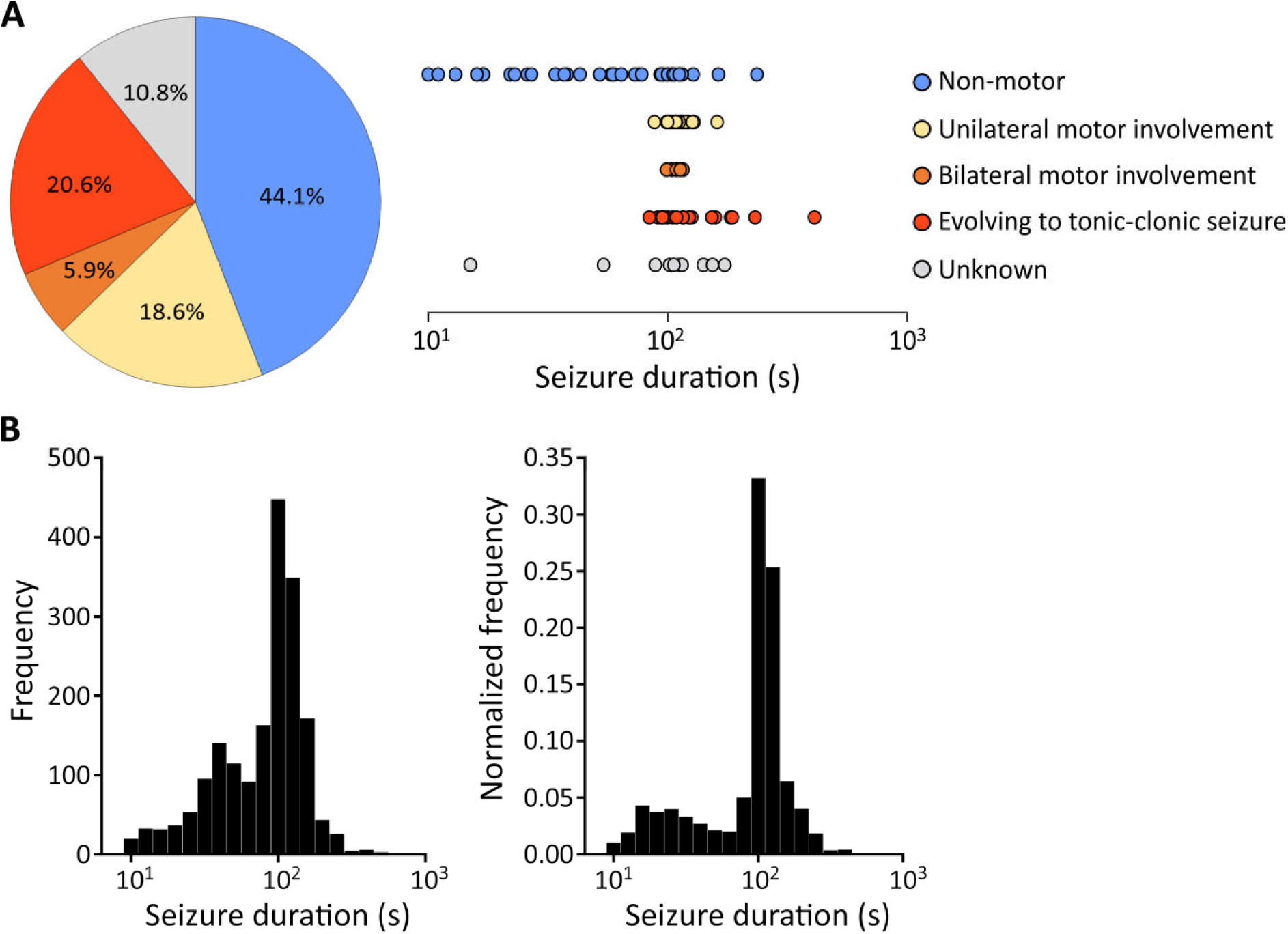
Behavioural manifestations of seizures. A, Semiology classification of seizures and its distribution of seizure duration (n = 102 seizures from 8 animals). The “unknown” means the behaviours were not visible because the animals were beneath the environmental material or interrupted video signals. B, Histogram of seizure duration from 10 animals over the whole recording period. Right: the frequency was normalized to the total seizures of individual animals.

### Epilepsy produced by TeNT in visual cortex presents as focal seizures with or without secondary generalization

A total of 102 randomly selected seizures (out of a total of 1717 seizures observed) from 8 epileptic rats were used to assess the correlated seizure behaviour by a neurologist (Figure 3A). 45 seizures (44.1%) were non-motor focal seizures presenting as behavioural changes (suppl. video 1), including repetitive eye blinking, sudden freezing or jumping, or agitated and aggressive searching behaviour which may be consistent with visual hallucinations arising from the epileptogenic focus. Nineteen seizures (18.6%) were classified as unilateral motor involvement, which manifested as contralateral limb twitching. Seizures propagating to bilateral motor symptoms were observed in a further 6 (5.9%) animals. Twenty-one seizures (20.6%) evolved to generalized tonic-clonic seizures (suppl. video 2). Apart from these observable seizures, associated behaviours were unobservable in 11 seizures (10.8%) due to the animals staying out of sight within the environmental enrichment material or due to transient interruption of video recordings. For non-motor seizures, the seizure duration ranged from very brief (10 s) to long seizures (> 100 s), whereas a relatively long and more consistent duration (∼100s) was typical for seizures with unilateral or bilateral motor involvements (Figure 3A, right panel). Seizures that evolved to tonic-clonic, activity were typically long, lasting from 200 s to 400 s.

The density distribution of seizure durations for all seizures from 10 animals are summarised in figure 3B (left panel). As the total number of seizures in two of the animals was extremely high relative to the remaining 8 animals, the density distribution was corrected by normalizing to total seizure counts for individual animals to avoid the two animals with highest frequency of seizures disproportionately affecting the distribution of seizure durations (Figure 3B, right panel). The bimodal pattern was shown where the highest probability of seizure duration was at 100 s to 125 s, and a smaller peak was at approximately 15 s to 25 s.

### No abnormal response to photic stimulation

As Sprague Dawley rats are thought to have poor vision^22^, and this may influence detection and provocation of seizures by visual stimulation, ability to trigger seizures in this model using intermittent photic stimulation was tested in TeNT-treated Lister Hooded epileptic rats (n = 3). However, none of the animals manifested photic-stimulation induced seizures, photoconvulsive (photoepileptiform) response or displayed prominent ECoG changes during photic stimulation.

### Seizures occurrence during light dark cycles

To determine whether there is a correlation between circadian rhythm and seizure occurrence, the distribution of seizures during day (light-on) and night (light-off) periods was analysed. The proportion of seizures occurring during different time periods was normalised within individual animals to control for the variability of seizure frequency between animals. For 7 of 10 animals, seizures predominantly occurred during the light periods (sleeping periods), one animal had equal seizure frequency during light and dark, and two animals had slightly higher seizure counts during the dark periods (active periods; Fig. S2). Overall seizure frequency gradually increased from the beginning of sleeping phase (7 a.m.), reaching the highest seizure activity during the mid-stage of sleep (1 p.m. – 4 p.m.), and slowly subsiding during the active dark period (7 p.m. – 7 a.m.) (Figure 4). There was no significant difference of seizure duration between day and night (Wilcoxon matched-pairs signed rank test, p = 0.275).

**Figure 4.**
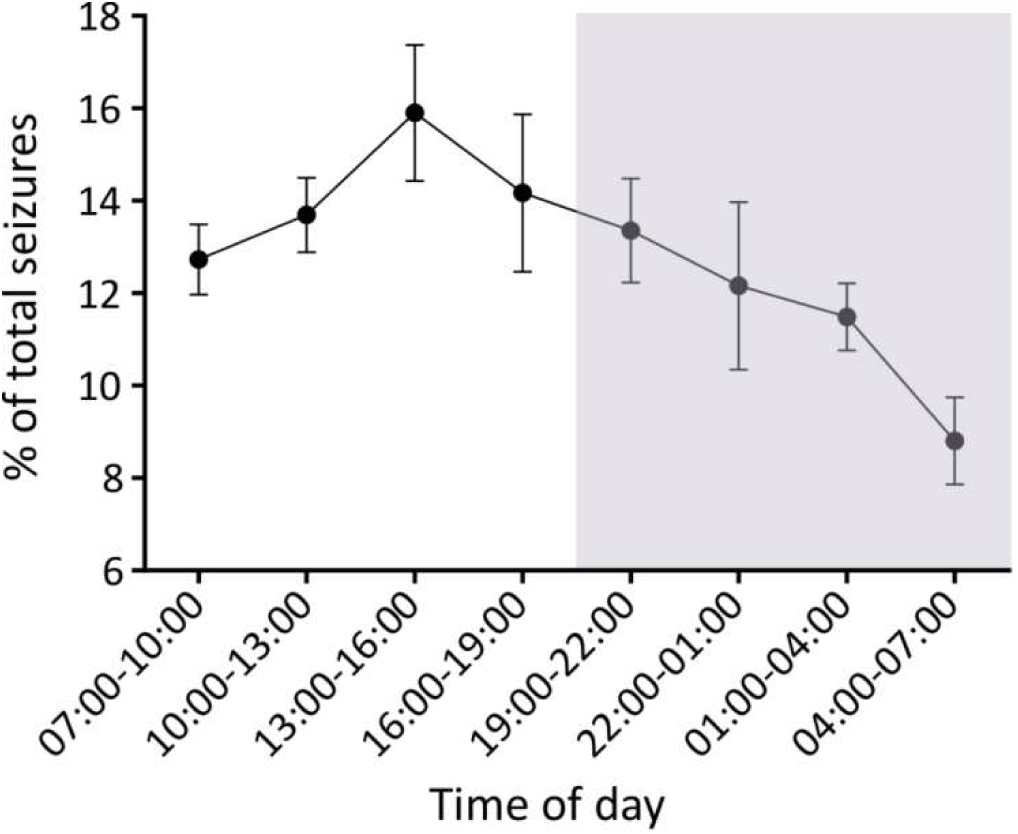
Seizure activity during the light/dark cycles. The grey shading represents the light-off/activity period (7 p.m. – 7 a.m.). Data are presented as mean ± SEM. (n = 10 animals)

### Clustering and periodicity of seizures

The distribution of seizures for individual animals over the recording period was not homogeneous (Figure 5A). The inter-seizure intervals were bimodally distributed with most of the ISIs between 3 to 4.5 hours (Figure 5B). However, no obvious correlation between seizure duration and ISIs was found (Fig. S3). To determine if the temporal distribution seen was a consequence of the dissimilar seizure rates between different animals, the ISI distributions for individual animals were compared to exponential distributions (Figure 5C). Indeed, most ISI distributions (7/10) differed significantly from exponential distributions, implying that seizure occurrence does not follow a homogeneous Poisson process, and not random.

**Figure 5.**
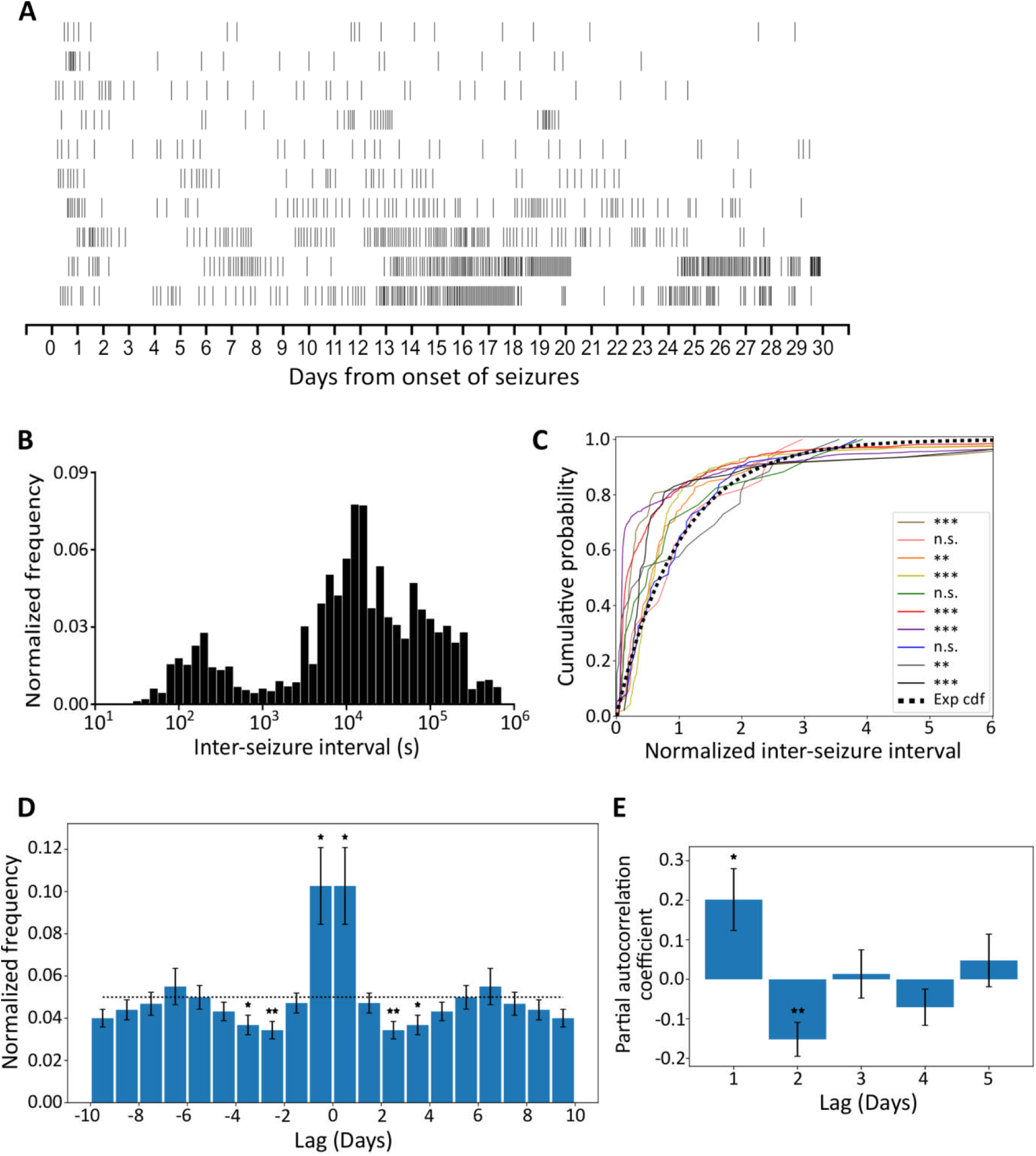
Seizures tend to occur in clusters and periodically. A, Raster plots of all seizures from onset of seizures over the whole recording period. B, Histogram of inter-seizure interval from 10 animals. The frequency was normalized to the total seizures of individual animals. C, Deviation between individual animals (color-coded) normalized inter-seizure-interval distributions and the exponential cumulative distribution function (Exp cdf). (Lilliefors-Komolgorov-Smirnov test, **p < 0.01, ***p < 0.001, n.s. = not significant). D, Autocorrelation function. Data are re-calibrated for short acquisition period (see methods in the main text) and presented as mean ± SEM. The dotted line represents the expected value of a uniform distribution. First, third and fourth day counts differ significantly from the uniform distribution (two-tailed one sample t-test, *p < 0.05, **p < 0.01). E, Partial autocorrelation coefficients for daily seizure counts (mean ± SEM). The first and second coefficients differ significantly from zero (two-tailed one sample t-test, *p < 0.05, **p < 0.01).

To characterize the temporal structure of seizure occurrence, we made a normalized peri-seizure histogram; this demonstrated that on average seizures tended to cluster (Figure 5D), i.e., there is a significantly higher probability of seizure occurrence if a seizure has occurred in the previous 24 hours (p = 0.02). This probability decreases to values significantly below the expected value for the period between 48 and 96 hours after seizure occurrence (p = 0.006 and p = 0.02). Partial autocorrelation analyses on daily seizure occurrence indicates that the seizures follow a periodic pattern (positive first coefficient, p = 0.03, and negative second coefficient, p = 0.008) (Figure 5E).

### An approach to refine the model by predicting the number of seizures from early data

There was a significant and notable correlation between the log number of seizures in 1st week (from onset of seizures) and the number that occurred during remaining recording period (day 8 – day 30; Figure 6A). We assessed this correlation for its predictive ability using a Gaussian process model with a linear kernel (Figure 6B, C and D). This model of the data indicates, for example, that if the number of seizures during the first week of seizures is between 11 and 34 for a given animal, that animal may be predicted to have between 20 to 100 more seizures in the following 23 days, with a confidence above 50%.

**Figure 6.**
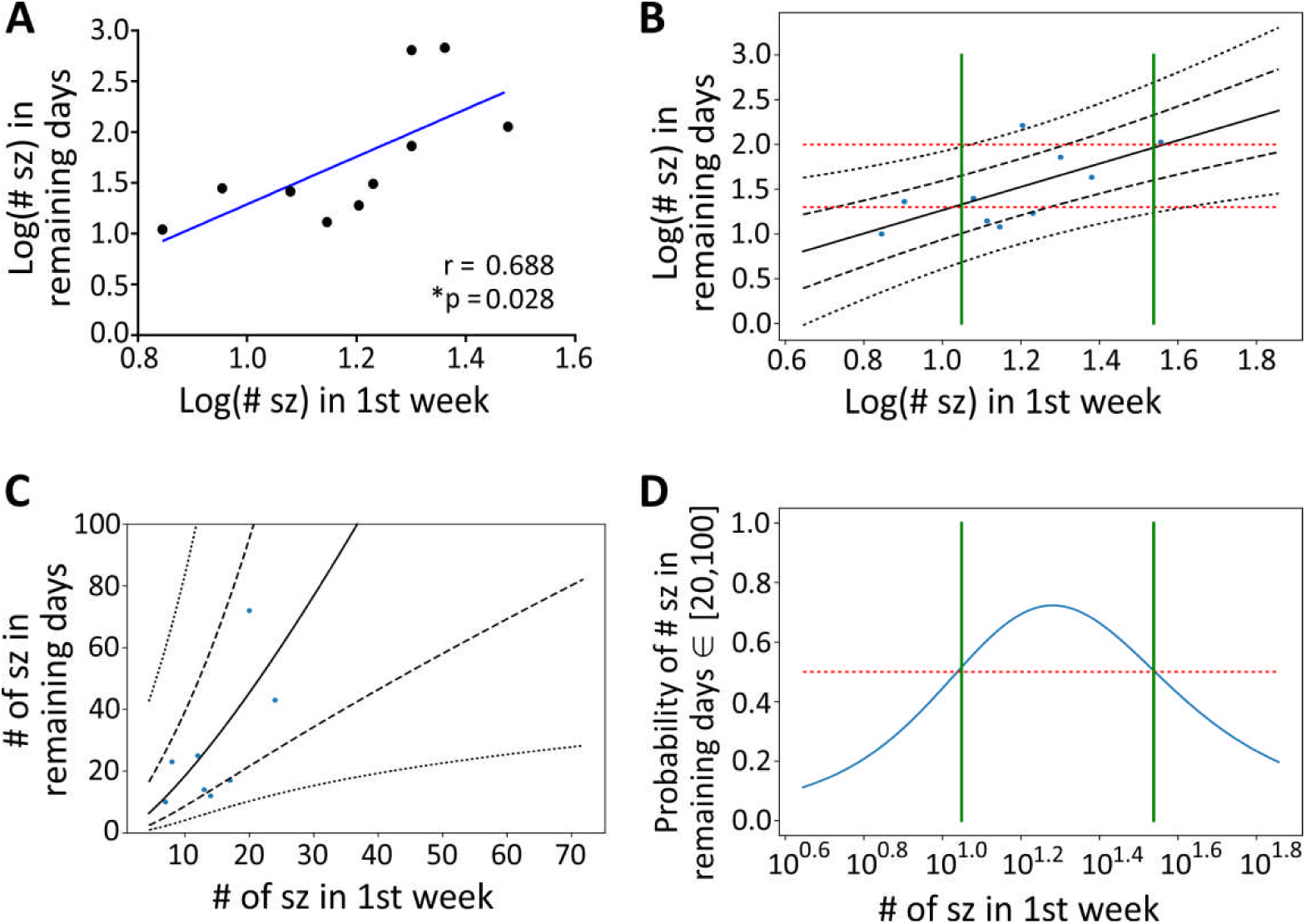
Predictions on the number of seizures. A, The plot of Pearson correlation coefficients for log number of seizures in first week against the number of seizures in days 8 to 30 after onset of seizures (n = 10 animals). B, Gaussian process modelling of the relation between the log number of seizures in first week after onset of seizures and the number of seizures in the remaining recording days. Black lines represent the posterior Gaussian distribution (mean ± one, and two SD). Red lines delimit a target interval of the number of seizures (from 20 to 100) in the remaining days and green lines delimit the interval for which there is more than 50% chance of observing such number of seizures (from 11 to 34). C, Same as B, but with the axes exponentially transformed. D, Probability that the number of seizures in the remaining days fall within the target interval of 20 to 100. Green lines delimit the interval above the 50% threshold mark (red line).

## Discussion

In this study, we demonstrate that application of TeNT into rat visual cortex produces a robust model of occipital lobe epilepsy. Although models of TeNT applied in mouse neocortex and rat motor cortex have been described,^13, 14^ this is the first time a chemically-induced focal neocortical epilepsy model exhibiting both clear ictal electrographic epileptic discharges and associated behavioural seizures which resemble human seizures. The high rate of induction of epilepsy associated with very low morbidity, mortality and no sudden unexpected death in epilepsy (SUDEP) indicates that this is a well-tolerated and reproducible rat model of epilepsy, and may be considered a refinement of other models (such as the tetanus toxin motor cortex model, where seizure activity can be accompanied by weight loss and death).^13^

The major advantages of the TeNT model of focal epilepsy are that epileptogenesis is not triggered by SE and does not rely upon neuronal loss or major disruption of tissue,^11^ and that it models focal neocortical epilepsy, which represents a major unmet need. TeNT is cleared from the brain a few days after local administration,^23^ but the disruption of VAMP persists, leading to recurrent spontaneous seizures over a long period.^14, 24^ In the visual cortex this non-SE induced epilepsy model has a relatively consistent latent period followed by progressively increasing seizure frequency and duration during the establishment of epilepsy which reaches a plateau, and then resolves.

We defined repetitive epileptic discharges lasting for more than 10 seconds as ictal activity based on most seizures seen in humans.^25–27^ This model exhibits both short (≤40 s) and long-lasting (>100 s) seizures with associated behavioural manifestations, and has markedly longer duration (usually 90 – 140 s) seizures than TeNT injection in motor cortex (which has mostly high frequency burst <1 s, or most seizures lasting only few seconds).^13^ This is probably due to higher doses (15 ng) injected into visual cortex being better tolerated than that injected into motor cortex (12.5ng).^13^ Despite the electrographic activity having only been recorded from an electrode in the epileptogenic zone, seizures propagated to and spread to other ipsilateral and/or contralateral brain regions as could be observed from the seizure behaviours. These characteristics resemble the majority of human focal epilepsies. A limitation of this model is that in some of the animals, the seizure frequency decreased between 4 to 5 weeks after the onset of spontaneous seizures. However, given the clinical need for new therapies and the paucity of robust focal neocortical models of chronic epilepsy, a model that recapitulates many features of human neocortical epilepsy is an invaluable tool for developing new treatments for focal neocortical epilepsy. Therefore, this model may be used to test for efficacy in developing new gene therapies and other novel treatment strategies.

Some specific human epilepsy syndromes, which encompass generalized and focal epilepsy, have potential links with photosensitivity, the propensity to trigger seizures by photic stimulation.^28, 29^ This is, however, rare in acquired occipital lobe epilepsies,^30, 31^ and similarly we were unable to trigger seizures with intermittent photic stimulation in our model. One explanation is that photosensitivity is not an intrinsic property of occipital lobe epilepsies but is largely an inheritable trait;^32^ indeed, it is possible to have a photoparoxysmal response without having epilepsy^33^ and gene mutations and polymorphisms have been identified that are strongly associated with photosensitivity.^34, 35^

Our data indicate that circadian cycles may affect seizure frequency, but not seizure duration. A major advantage of using experimental models over human studies is that they are not confounded by treatment or other environmental factors can be controlled. We found a trend towards higher seizure frequency in light periods which is consistent with a lower seizure threshold in the sleeping state. The epileptogenic process may had an impact on the reciprocal influences between sleep-wake cycles and epilepsy.^36^ Higher activity of seizures and interictal epileptiform discharges during non-rapid eye movement (NREM) sleep has been shown in some human epilepsies and in some experimental epilepsy models.^37, 38^ Approximately 20% of people with epilepsy have predominately nocturnal seizures.^39^ Thalamocortical synchronization has been invoked to account for this association.^40, 41^ Circadian changes at many different levels have been proposed to explain the relationships between epilepsy and circadian rhythm.^36, 42^ Further studies to explore dynamic molecular changes regulating the interactions of circadian rhythmicity, epileptogenesis and seizure occurrence is important to understand the underlying mechanisms.

Clustering of seizures is often seen in human epilepsies.^43^ In this model, multiple seizures had a higher probability of occurring within a day, and the overall seizure occurrence exhibited periodicity in most of the animals. Seizure clustering could be due to positive feedback mechanisms, whereas the relative refractoriness after a cluster might be caused by slower negative feedback mechanisms.^44^ The rhythmic phenomenon of seizure activity may be exploited to refine treatments to prevent occurrence of following seizures.

The total number of seizures experienced among individual animals in this model was highly variable. However, a positive correlation between the number of seizures in the first week and the remaining recording period permits a seizure prediction algorithm which can be applied to control for the variability of seizure frequency in animals enrolled in studies comparing treatments.

In summary, an optimized rat model of visual cortical epilepsy mimics focal neocortical epilepsy in humans, providing a non-SE and non-lesion induced epilepsy model for studying human epilepsy and seeking novel therapeutic strategies. We further provide a model for predicting number of seizures, which will be valuable to reduce the effect of variability of the model on the interpretation of therapeutic interventions.

## Acknowledgements

We thank G. Schiavo (UCL) for the gift of tetanus toxin, and Lenon Gwaunza for early analysis of this model. This work was supported by the Medical Research Council, the Royal Society, the Wellcome Trust, Epilepsy Research UK, and the Chang Gung Memorial Hospital, Taiwan (CMRPG3C1021, CMRPG3C1022, CMRPG3C1023, CMRPG3C1024, and CMRPG3C1025).

## Disclosure

We confirm that we have read the Journal’s position on issues involved in ethical publication and affirm that this report is consistent with those guidelines. None of the authors has any conflict of interest to disclose.

