## Supplementary Materials for "Semiology, clustering, periodicity and natural history of seizures in an experimental visual cortical epilepsy model"

Video 1: non-motor seizure

Video 2: focal seizure evolving to generalized tonic-clonic seizure

Available from authors upon request.

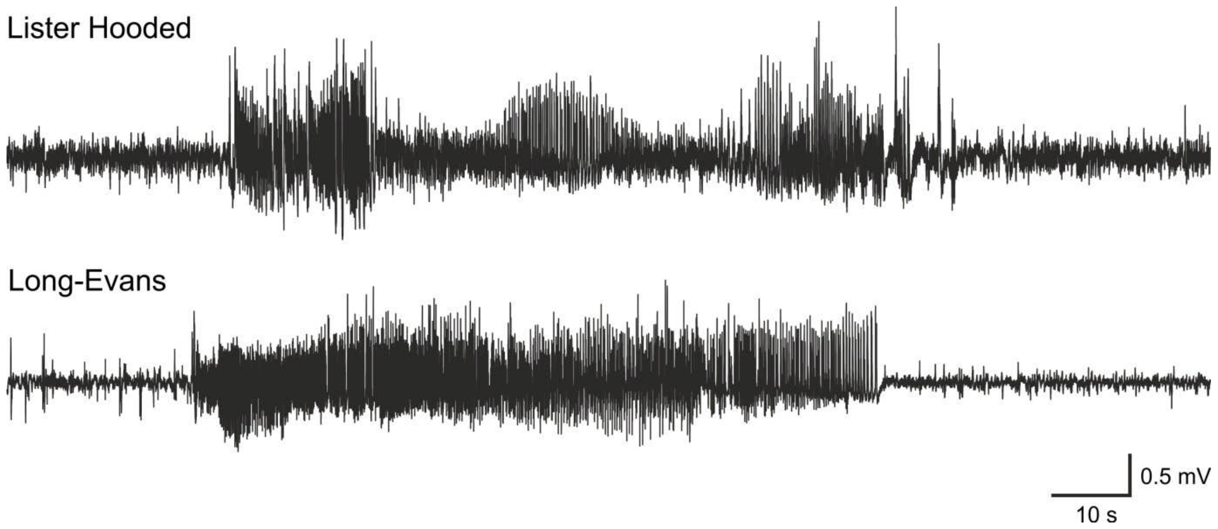


**Figure S1** Representative seizures from Lister Hooded and Long-Evans rats. The TeNT model of visual cortical epilepsy can be induced in other rat strains.

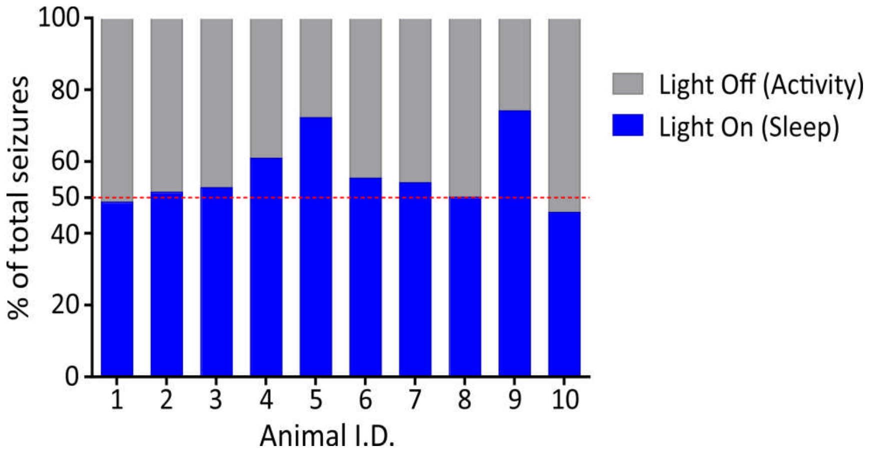


**Figure S2** Distribution of seizure occurrence during day and night for individual animals. There are higher proportions of seizures during sleeping period in most of the animals.

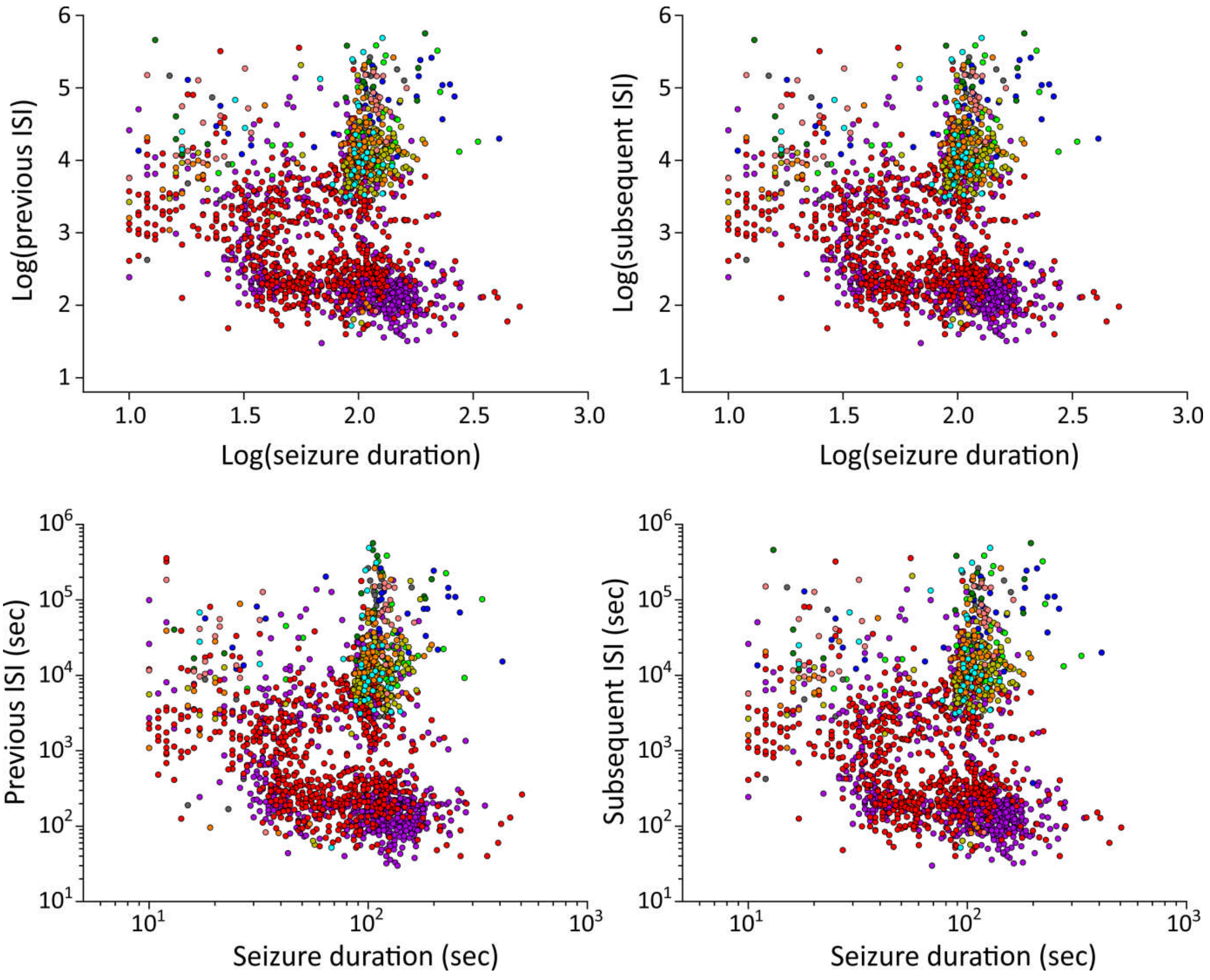


**Figure S3** No obvious correlation between seizure duration and ISI. Upper: Scatter plot of log of seizure duration with log of previous (left) and subsequent (right) ISI. Lower: Scatter plot of seizure duration with previous (left) and subsequent (right) ISI.
